# Lineage-specific positive selection signatures identify stress-response genes in wild wheat relatives

**DOI:** 10.64898/2026.09.22.740609

**Authors:** Manuel Barrientos Tecún, Vincent Ranwez, Nathalie Chantret, Concetta Burgarella

## Abstract

Understanding how crop wild relatives (CWR) adapt to diverse environments is important both for evolutionary biology and for the development of climate-resilient agriculture. CWR represent an important reservoir of adaptive variation that can be exploited for future crop improvement. To investigate how natural selection has shaped adaptation in these species, we analysed whole-transcriptome data from four diploid wild wheat relatives with contrasting mating systems: the outcrossing species *Aegilops speltoides* and *Aegilops mutica*, and the selfing species *Aegilops tauschii* and *Triticum urartu*. We combined the imputed McDonald–Kreitman test (MKT) with lineage-specific estimates of divergence obtained from branch-specific models of molecular evolution. This framework reduces biases caused by slightly deleterious mutations and enables adaptive substitutions to be assigned to individual evolutionary lineages. We detected marked differences among species in the prevalence of positive selection. Evidence of adaptive evolution was concentrated in the outcrossing species, whereas few positively selected genes were identified in the selfing species. Comparisons between standard and branch-specific MKT approaches showed that some signals initially inferred as shared among species were reassigned to individual lineages when divergence was estimated on a branch-specific basis, emphasizing the importance of accurately localizing adaptive substitutions. Functional annotation revealed several candidate genes containing domains associated with abiotic and biotic stress responses. Our study provides a framework for detecting positive selection in wild wheat relatives and identifies candidate genes that may contribute to adaptation and represent valuable targets for future functional analyses and wheat improvement programs.

## 1 Introduction

Global food security is increasingly threatened by climate change and population growth. Climate change reduces crop yields, increases yield instability, and increases food prices through more frequent heat waves and droughts (*Climate change 2021* 2021). The expanding range of pests, weeds, and pathogens further threatens agricultural productivity (Bajwa et al. 2020). Genetic diversity within crops is a key resource to cope with these challenges because it increases the likelihood that crops possess alleles conferring tolerance to abiotic stresses and resistance to emerging pests and diseases (Henry and Nevo 2014; H. Zhang, Mittal, et al. 2017; Dempewolf et al. 2017; “Utilizing Crop Wild Relatives to Combat Global Warming” 2019; Cockel et al. 2022).

However, this adaptive potential has been strongly reduced in modern crops due to domestication and breeding bottlenecks. Early assessments by the Food and Agriculture Organization of the United Nations (FAO) indicated that by the late twentieth century, approximately three-quarters of the genetic diversity present in traditional crop varieties had already disappeared from cultivated fields (Food and Agriculture Organization of the United Nations 1993). More recent syntheses with large-scale analysis show that more than 95% of studies report changes in crop genetic diversity, and nearly 80% provide evidence for a net loss (Khoury et al. 2022). This reduced genetic diversity limits stress adaptation and increases vulnerability to pests and climate change (Dempewolf et al. 2017; Langridge et al. 2022).

To broaden the genetic base of modern crops, crop wild relatives (CWR) are a promising yet underutilized resource. These species often thrive in extreme environments and, therefore, have a vast reservoir of genetic variation for traits critical to agricultural sustainability, including tolerance to abiotic stresses and resistance to pests and diseases (Dempewolf et al. 2017; Brozynska et al. 2016; H. Zhang, Mittal, et al. 2017). Several resistance genes have already been successfully transferred from CWR to cultivated species, demonstrating their considerable agronomic value (Farooq et al. 2025; Y. Li et al. 2025). Nevertheless, only a small fraction of the genetic potential of CWR has been exploited (Ford-Lloyd et al. 2011; Dempewolf et al. 2017; Leigh et al. 2022). If such adaptive traits can be identified and mobilized more systematically, they could contribute significantly to improving crop resilience and ensuring future food security.

QTL mapping and GWAS approaches have proven effective in identifying genomic regions and alleles associated with agronomically important traits in CWR (Khan et al. 2021; Tirnaz et al. 2022; Gojon et al. 2023) however, these methods rely on known phenotypes and trait variation. By contrast, analyzing patterns of adaptive molecular evolution in genomes and transcriptomes allows us to detect signatures of positive selection directly from sequence data. Such signatures indicate regions that have been functionally important during past evolutionary history and may harbor variants that contribute to stress resilience even in the absence of prior phenotypic characterization (Henry and Nevo 2014). This sequence-based perspective not only complements trait-focused studies, but also offers a systematic framework to rapidly uncover novel candidate alleles for wheat improvement (Afzal et al. 2019; L. Gao et al. 2023; Sthapit et al. 2024).

Numerous statistical methods have been developed to analyze genomic data and detect molecular adaptation across different evolutionary timescales. Divergence-based approaches compare nonsynonymous and synonymous substitution rates between species or along phylogenies to identify lineage-specific accelerations in protein evolution (Z. Yang 2007; Sahm et al. 2017; Álvarez-Carretero et al. 2023). In addition, polymorphism-based methods use intraspecific genetic variation, with statistics such as Tajima’s D (Tajima 1989), Fu and Li’s D (Fu and W. H. Li 1993) and haplotype-based tests (Kirsch-Gerweck et al. 2023) detecting selection from allele frequency patterns and linkage disequilibrium. However, both approaches face challenges, as signals of selection can be confounded by demographic history or population structure.

To address this, methods that jointly analyze polymorphism and divergence provide a more robust framework for disentangling adaptation from neutral evolutionary processes. Among these approaches, the McDonald–Kreitman test (MKT) (McDonald and Kreitman 1991) is a widely used method to detect positive selection in protein-coding genes. Unlike divergence-based approaches that rely on the ratio *ω*, which compares the number of synonymous (D_s_) and nonsynonymous (D_N_) substitutions, the MKT integrates both divergence (D_s_, D_N_) and polymorphism (P_s_, P_N_) data. Under neutrality, the ratio of nonsynonymous to synonymous changes is expected to be similar for polymorphic and fixed sites. Deviations from this expectation provide evidence for selection: an excess of nonsynonymous divergence relative to polymorphism indicates recurrent positive selection, whereas an excess of nonsynonymous polymorphism reflects the segregation of slightly deleterious mutations (SDMs). The MKT, by incorporating polymorphism data, accounts for purifying selection and has more power compared to divergence-only methods. However, MKT assumes neutrality at segregating sites, yet multiple species display an excess of SDMs (Smith and Eyre-Walker 2002; Messer and Petrov 2013; Galtier 2016). SDMs persist at low frequencies due to purifying selection and are rarely fixed. Consequently, they contribute to nonsynonymous polymorphism, leading to an underestimate of the proportion of adaptive substitutions (Eyre-Walker and Keightley 2009; Galtier 2016; Murga-Moreno, Coronado-Zamora, Casillas, et al. 2022). Additionally, The MKT covers the timescale from the present back to the divergence with an outgroup, so it cannot assign adaptive substitutions to specific lineages. Consequently, many studies have relied on divergence-based phylogenetic methods that do not incorporate polymorphism data to detect lineage-specific selection signatures.

To overcome these limitations, several extensions of the classical MKT framework have been developed at the gene level (Fay et al. 2001; Begun et al. 2007; Mackay et al. 2012; Murga-Moreno, Coronado-Zamora, Hervas, et al. 2019; Murga-Moreno, Coronado-Zamora, Casillas, et al. 2022). A recent version, the imputed MKT (impMKT) (Murga-Moreno, Coronado-Zamora, Casillas, et al. 2022) provides a robust approach using the observed site frequency spectrum to estimate and correct the proportion of SDMs contributing to nonsynonymous polymorphism. This strategy improves the accuracy of adaptive substitution rate estimates, even in the presence of demographic fluctuations and weak purifying selection.

In this study, we combine different MKT-based approaches with branch-specific divergence estimates obtained using PAML (Z. Yang 2007) that allow the assignment of adaptive substitutions to specific lineages while accounting for SDMs. We apply this framework to species from the Triticeae tribe, a major lineage within the grass family (Poaceae), which includes some of the world’s most important cereal crops such as wheat, barley and rye. Although Triticeae comprise approximately 500 species, the extent of adaptive genetic variation within this group remains largely unexplored (Leigh et al. 2022).

To identify adaptive variation of potential agronomic relevance, we focus on four diploid species from the Triticeae tribe: *Triticum urartu*, *Aegilops tauschii*, *Aegilops speltoides* and *Aegilops mutica* Figure 1. These species are closely related to cultivated wheat, with *T. urartu* and *Ae. tauschii* are donors of the A and D subgenome of hexaploid wheat, while *Ae. speltoides* is related to the B subgenome. *Ae. mutica* plays a central role in the D lineage through ancient hybridization (Glémin, Scornavacca, et al. 2019). These species also differ in mating systems: *Ae. speltoides* and *Ae. mutica* are outcrossing, while *Ae. tauschii* and *T. urartu* are largely selfing (Burgarella et al. 2024). Moreover, they occupy a wide range of ecological niches in the Fertile Crescent and adjacent regions (Zohary et al. 1969; Johnson 1975; Ohta and Saruhashi 2004; Hodgkin et al. 2008; J. Wang et al. 2013; Feldman and Levy 2023) o we can expect that they harbor diverse adaptations.

**Figure 1:**
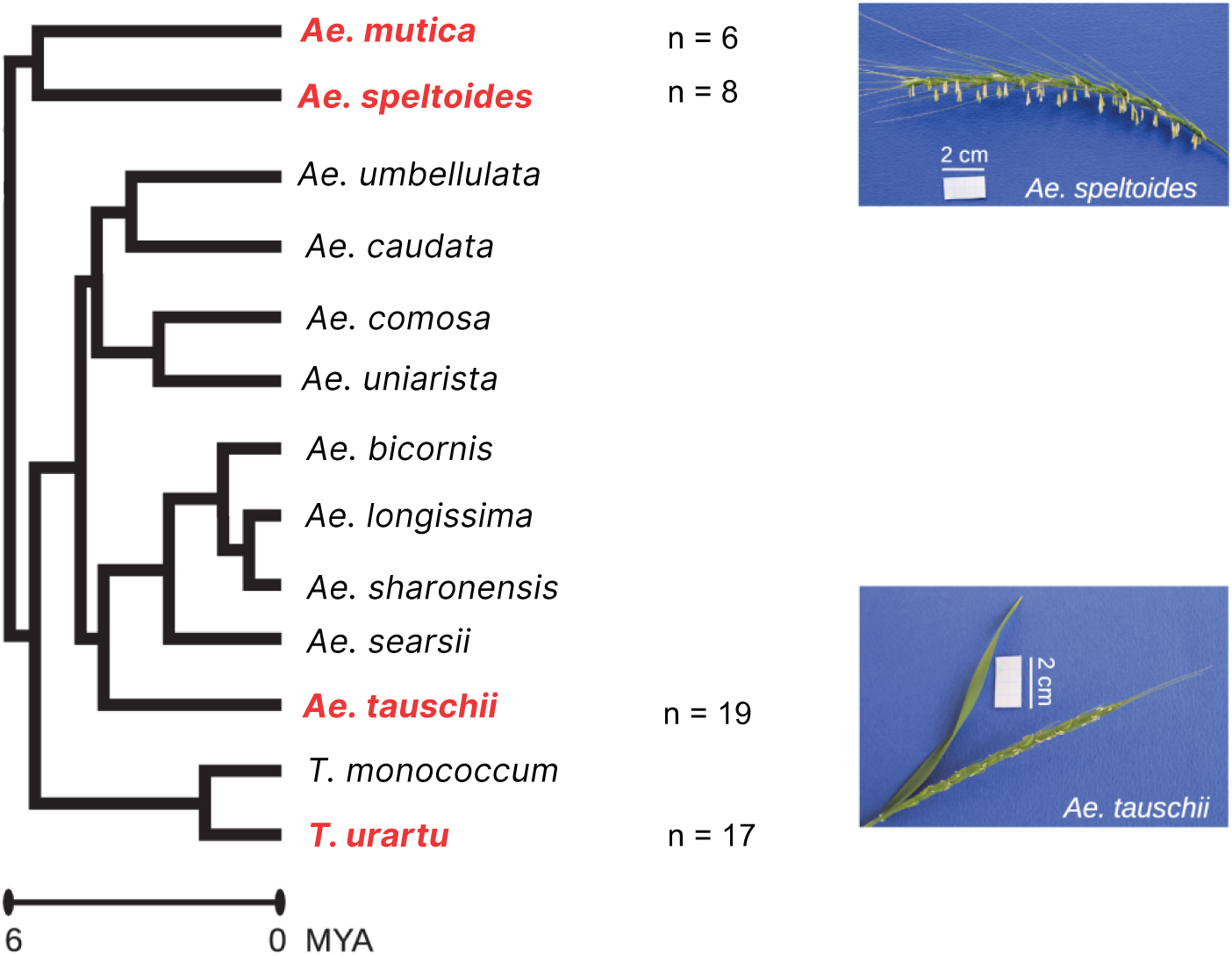
Phylogenetic relationships of diploid wheat species. Phylogenetic tree of the 13 diploid species within the genus, including the number of individuals available per species (adapted from (Burgarella et al. 2024)). Species names in red are the focal species used in this analysis. *Ae. speltoides* and *Ae. mutica* are known to be self-incompatible, while *Ae. tauschii* and *Triticum* sp. are predominantly self-fertilising (highly selfing). The remaining species are self-compatible.

Taking advantage of this combination of phylogenetic proximity, our approach aims to (i) detect adaptive evolution at the gene level, (ii) assign these signatures to specific evolutionary lineages, and (iii) characterize the functional categories of genes under selection. Using this framework, we identified several candidate genes under positive selection in wild wheat relatives (WWR) and inferred their lineage of origin. Functional annotation further reveals that many of these genes are associated with responses to biotic and abiotic stresses. This framework can be used as a tool to prioritize genetic variants from CWRs, facilitating their targeted incorporation into wheat improvement programs.

## 2 Methods

### 2.1 Dataset

Our analysis focuses on 4 diploid species of the genus *Aegilops/Triticum* (Triticeae tribe), the two self-incompatible (SI) outcrossing *Ae. speltoides* and *Ae. mutica* species, and two predominantly selfing *Ae. tauschii* and *T. urartu* species. In total 50 individuals were analyzed, 6 individuals for *Ae. mutica*, 8 individuals for *Ae. speltoides*, 19 individuals for *Ae. tauschii* and 17 individuals for *T. urartu*, (Figure 1), This data set is part of the data set of (Glémin, Scornavacca, et al. 2019; Burgarella et al. 2024). *Ae. speltoides* and *Ae. mutica* served as reciprocal outgroups for each other; *Ae. longissima* was used as and outgroup for *Ae. tauschii* ; and *T. monococcum* was used as an outgroup for *T. urartu*.

### 2.2 Sequencing and Transcriptome Assembly

RNA sequencing data were generated in previous studies (Sarah et al. 2017; Glémin, Scornavacca, et al. 2019; Burgarella et al. 2024). Briefly, RNA was extracted from mixed tissues (20% leaves and 80% inflorescences) of plants grown under the same conditions and used to construct TruSeq stranded mRNA libraries. Sequencing was performed on an Illumina HiSeq3000 platform (Get-PlaGe, INRAE Toulouse), producing paired-end RNA-seq reads. For the present study, we used the cleaned paired-end reads (FASTQ files) from the dataset deposited by Burgarella et al. (2024) in the Sequence Read Archive (SRA) under project PRJNA945064 (submission number SUB12943046).

### 2.3 Mapping and Coverage Analysis, Genotype Calling

For each focal species, we constructed a reference transcriptome by extracting annotated mRNA sequences from a publicly available reference genome: *Ae. speltoides* (L.-F. Li et al. 2022), *Ae. mutica* (Grewal et al. 2025), *Ae. tauschii* (Luo et al. 2017), and *T. urartu* (Ling et al. 2018). For outgroup species, we used *Ae. longissima* (Lux 2022) and *T. monococcum* (Ahmed et al. 2023). From each genome, we took the Ensembl canonical transcripts when available or highly confident mRNAs otherwise; for genes with several transcripts where neither was available, we selected the longest one (Table 1).

**Table 1:** Summary of transcriptome filtering and filtering statistics for the four focal species.

| Focal species | Retained mRNA | Well covered genes | Alignments with ortholog | Analyzable genes |
| --- | --- | --- | --- | --- |
| *Ae. speltoides* SI | 37,209 | 9,611 (25.82%) | 7,952 (21.37%) | 5,336 (14.34%) |
| *Ae. mutica* SI | 40,905 | 10,264 (25.09%) | 7,465 (18.24%) | 4,483 (10.95%) |
| *Ae. tauschii* HS | 39,597 | 8,656 (21.86%) | 7,137 (18.02%) | 2,626 (6.63%) |
| *T. urartu* HS | 37,961 | 9,996 (26.33%) | 5,967 (15.71%) | 975 (2.56%) |

“Retained mRNA” corresponds to the number of reference transcripts kept after filtering. “Well-covered genes” refers to genes with at least 50% of positions covered ( $\geq 10\times$ depth in $\geq 6$ individuals). “Alignments with ortholog” indicates the number of genes successfully aligned with the outgroup orthologous sequence. “Analyzable genes” are those that passed quality and polymorphism filters (minimum coding length, at least 5 polymorphic sites and one polymorphic and divergent site for both synonymous and nonsynonymous categories) and are thus eligible for selection analyses. Percentages in parentheses are relative to the total number of retained mRNA for each species. SI: self-incompatible (outcrossing); HS: highly selfing (autogamous).

We then used VESPA (Webb et al. 2017) to retain only protein-coding sequences with complete open reading frames—i.e., sequences divisible by three and containing no premature stop codons. After these steps, we extracted mRNA sequences including 200 bp upstream and downstream flanking regions, from their respective genome assemblies using AGAT (Dainat et al. 2024). The final filtered reference transcriptomes were used as mapping targets. The mappings were performed using Gecko workflows (Ardisson et al. 2024). Because soft-clipped read clusters often indicate structural variants (Yan et al. 2021) or ambiguous mapping regions, we removed these reads from our data set. After mapping, we defined “covered” positions as genomic sites with at least 10 reads in a minimum of 6 individuals. Genes were classified as “well-covered” if at least 50% of their positions met this coverage threshold and only these high-quality genes were retained for subsequent analyzes.

From the BAM files, a multi-FASTA was generated using Reads2SNP v2.0 **reads2snp**(http://kimura.univ-montp2.fr/PopPhyl/resources/tools/reads2snp.tar.gz). This tool is specifically designed to analyze transcriptome data for population genomics of nonmodel species. The method first calculates the posterior probability of each possible genotype in the maximum-likelihood framework after estimating the sequencing error rate. Genotypes supported with a probability higher than a given threshold (here 0.95) are retained; otherwise missing data are called. We required a minimum coverage of 10× per position and per individual to call a genotype. SNPs are then filtered for possible hidden paralogs (duplicated genes) using a likelihood ratio test based on explicit modeling of paralogy (“paraclean” option embedded in the reads2snps software. Because genotype calling depends on the expected level of heterozygosity, we used the inbreeding coefficients (*F*_is_) from (Burgarella et al. 2024): 0.36 for *Ae. speltoides*, 0.18 for *Ae. mutica*, 0.93 for *Ae. tauschii*, and 0.90 for *T. urartu*.

### 2.4 MKT analysis

To test for evidence of positive selection, we individually analyzed genes using two MKT frameworks. The first, which we refer to as the two-species MKT, corresponds to the standard MKT framework and relies on a single outgroup species. Within this framework, we used the standard MKT (McDonald and Kreitman 1991) and the impMKT (Murga-Moreno, Coronado-Zamora, Casillas, et al. 2022). The second framework, which we refer to as the three-species MKT, uses two outgroups species. In this framework, we applied a branch-specific version of the MKT (Figure 2) and impMKT, allowing the assignment of substitutions to the focal lineage. For comparison, we also performed the MKT and impMKT in the context of three-species (i.e., using the same alignments but with only one outgroup).

**Figure 2:**
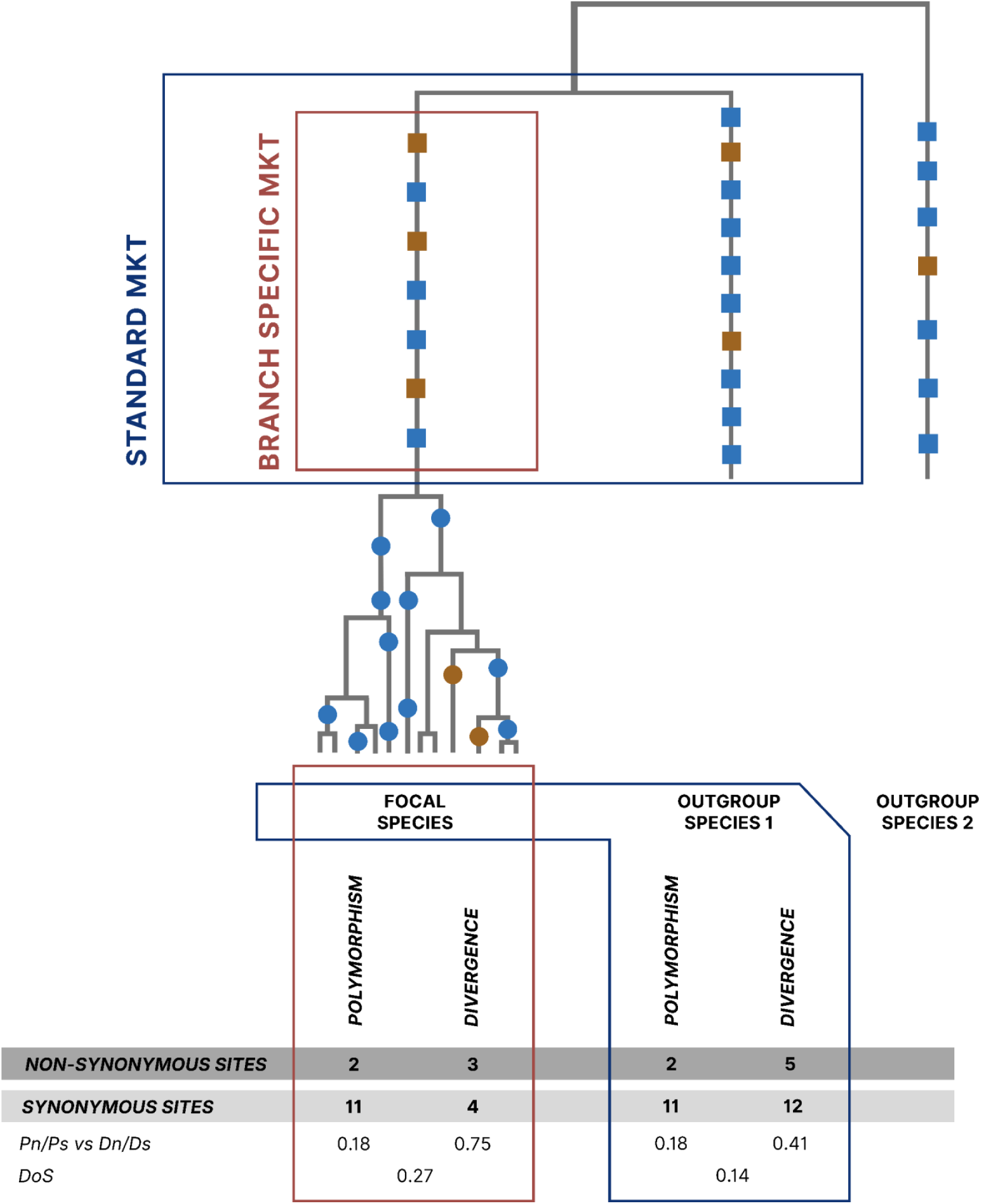
Visualization of polymorphism and divergence for an MKT. An MKT contrasts the ratio of changes of nonsnonymous to synonymous sites for polymorphisms of a focal species (*Pn/Ps*) with fixed differences between species (*Dn/Ds*). Synonymous sites are used as a neutral reference to test for selection on nonsnonymous sites. The genealogies in the diagram depict non-synonymous changes as red dots and synonymous changes as purple dots for polymorphism, and nonsynonymous changes as red squares and synonymous changes as purple squares for divergence. In a standard MKT, divergence is estimated from fixed differences between the focal species and a single outgroup, such that substitutions accumulated along both lineages contribute to *Dn* and *Ds*. Consequently, an excess of non-synonymous divergence cannot be unambiguously attributed to either lineage. By including a second outgroup, substitutions can be polarized and divergence can be estimated specifically along the focal lineage, allowing adaptive evolution to be inferred for the focal species alone.

#### 2.4.1 Polymorphism Estimates

For both two-species MKT and three-species MKT frameworks we applied the same method to obtain polymorphism estimates. The CDS sequences were aligned using MACSE v2.07 with alignSequences program (Multiple Alignment of Coding Sequences) (Ranwez et al. 2021). Polymorphism counts and stats for each gene alignment were obtained with EggLib version 3.5.2 (Siol et al. 2022), using a threshold of 30% of missing values allowed. For selfing species *Ae. tauschii* and *T. urartu*, all statistics were calculated on n/2 alleles by randomly drawing one haploid sequence per gene and individual.

We used a 15% frequency threshold to separate effectively neutral variants (above) from SDM (below). The neutral expectation is 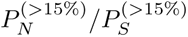, and the number of SDMs was estimated as:

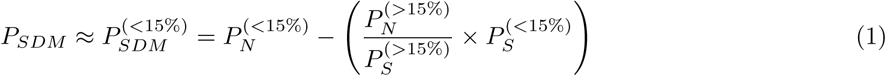

#### 2.4.2 Divergence Estimates

#### 2.4.3 Two-species MKT

To run the standard and impMK tests in the two species framework MKT, we estimate divergence at synonymous and nonsynonymous sites between each focal species and an outgroup species. We identified orthologous genes through BLAST analysis using a Reciprocal Best BLAST (RBB) approach between the CDS sets of the reference species and the outgroup species.

We created multi-FASTA files per gene with polymorphism sequences of the focal species and the outgroup sequence. Prior to alignment, non-homologous regions were removed from all multi-FASTA files using the TrimNonHomologousFragments option in MACSE, and coding sequences were aligned with alignSequences. Genes containing frameshift codons in initial alignments were realigned with increased frameshift penalties (-fs 320 -fs_term 320) to reduce spurious frameshift calls. Only CDS alignments free of frameshift symbols were retained. Counts of *Dn* and *Ds* were calculated with dNdSpNpS v.1.0 (available at https://kimura.univ-montp2.fr/PopPhyl/index.php?section=tools), which relies on the Bio++ libraries (Dutheil et al. 2006), with default parameters values.

#### 2.4.4 Three-species MKT

To estimate branch-specific divergence for each focal species, we constructed species triplets that include the focal species and two closely related species. This approach allows us to define a phylogenetic tree and estimate the divergence specifically on the branch of the focal species (Figure 2). The species triplets used for each focal species are shown in the (Table 3), orthologous genes were identified using OrthoFinder v3.0.1b1 (Emms and Kelly 2019).

Orthologous CDS were first aligned at the amino-acid level with MAFFT (v7.515) (Katoh and Standley 2013), and alignments were cleaned for non-homologous segments using HMMcleaner (Di Franco et al. 2019). We quantified per-sequence coverage as the fraction of alignment columns that contain a canonical nucleotide (A, C, G or T) in that sequence. Positions corresponding to gaps (“–” or “.”), Ns or any other ambiguous character were treated as uncovered. Only alignments in which all sequences showed at least 80% coverage were retained. As codeml takes nucleotide alignments as input, we translated all protein alignment into nucleotides with pal2nal (v14.1) (Suyama et al. 2006).

To ensure that polymorphism and divergence were estimated from the same set of codon sites, we next combined, for each focal species, the within-species CDS alignment (all individuals of that species) with the corresponding multi-species ortholog alignment. This was done with the enrichAlignment program in MACSE, which adds the reference and outgroup sequences to the polymorphism alignment while preserving the codon frame. On the resulting enriched alignments, we masked codon columns where (i) the outgroup sequence was missing, or (ii) fewer than six sequences were present in the focal species (after accounting for missing data). Masked codons were replaced by “NNN” in the nucleotide alignment. We then computed per-sequence coverage as above and kept only enriched alignments whose mean coverage across sequences was at least 70%, so that both polymorphism and divergence statistics were based on well-covered, homologous codon sites.

The divergence used in the branch-specific MKT was obtained with the CODEML software, part of the PAML package (Z. Yang 2007) using the branch model that assumes different *ω* ratio parameters for different branches in the phylogeny, allowing estimation of 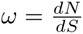 in the branch of the focal species. For each gene, we converted the rates *dN* and *dS* to counts of divergent sites by multiplying *dN* and *dS* by the corresponding numbers of nonsynonymous and synonymous sites, respectively, and rounding the resulting values to the nearest integer to obtain counts of *Dn* and *Ds*. For comparison, standard MKT and impMKT analyzes were conducted on the same set of gene alignments, restricting the data set to a single outgroup.

### 2.5 Statistical significance

We computed all versions of MKT on genes that satisfied the following criteria: (i) at least five polymorphic sites in total (*Pn* + *Ps ≥* 5), (ii) at least one site in each of the four MKT categories (*Pn*, *Ps*, *Dn*, *Ds*), *Dn* and *Ds* represent the numbers of nonsynonymous and synonymous substitutions per gene, respectively, and *Pn* and *Ps* represent the numbers of nonsynonymous and synonymous polymorphisms. For the impMKT and branch specific impMKT version, we used the same filters after correcting the non-synonymous polymorphism count by subtracting the estimated number of SDMs from the *Pn* category before computing the test. To test whether the ratio of nonsynonymous to synonymous changes differs significantly between polymorphism and divergence, we performed Fisher’s exact test on the contingency table of polymorphism and divergence counts. To further quantify deviations from neutrality and to distinguish whether a gene shows an excess of *Pn* or an excess of *Dn* we computed for all genes the direction of selection (*DoS*) as in (Stoletzki and Eyre-Walker 2011):

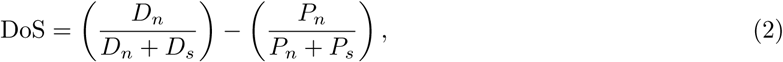

For each MKT version, genes were considered to show evidence of positive selection if they exhibited a DoS *>* 0 and a *p*-value *<* 0.05. These genes are hereafter referred to as *positively selected genes*. For gene-level analyzes (functional annotation, gene expression, protein–protein interaction networks, accelerated protein evolution, and recombination rate associations), we defined a subset of *candidate genes* as those identified as positively selected by either the branch-specific MKT or the branch-specific impMKT approach, or both.

### 2.6 Association Between adaptative evolution and Signatures of Accelerated Protein Evolution

To determine whether candidate genes show signatures of accelerated sequence evolution, we used CODEML branch-model likelihood-ratio tests. For each gene, we compared the branch-model *ω* ratio (allowing a distinct *ω* on the focal lineage) with the one-ratio model (M0) using:

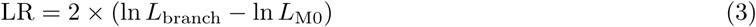

Genes with a significant LRT and a foreground *ω* greater than the background *ω* were classified as branch-model accelerated. The overlap between these accelerated genes and our candidates was quantified and enrichment was tested using Fisher’s exact tests.

### 2.7 Expression Analysis: Abiotic and Biotic Stress of Candidate Genes Under Positive Selection

Stress-responsive expression data were obtained from (Ramírez-González et al. 2018) which provides RNA-seq profiles of *Tr. aestivum* exposed to multiple abiotic and biotic stress conditions. Orthologs in *T. aestivum* were identified for all genes analyzed by branch-specific MKT/impMKT methods in *Ae. mutica* and *Ae. speltoides*. We then compared the stress-responsive expression profiles of candidate genes identified by the selection analyzes with those of non-candidate genes in several stress conditions, including heat stress (Z. Liu et al. 2015), cold stress (Seifert et al. 2016), phosphate starvation (Oono et al. 2013), powdery mildew infection (H. Zhang, Y. Yang, et al. 2014), fusarium infection (Schweiger et al. 2016), and septoria infection (Rudd et al. 2015).

Differential expression analysis was conducted using the DESeq2 package in R (Love et al. 2014), which models RNA-seq count data with a generalized negative binomial linear model (GLM). The experimental design was defined as a condition, allowing pairwise comparisons of each stress treatment against the control. Genes were considered significantly differentially expressed when exhibiting an adjusted p-value (padj < 0.05). To focus on biologically significant changes in expression, genes were classified as upregulated when they showed padj < 0.05 and a log_2_ fold change greater than 0.5 and downregulated when padj < 0.05 and a log_2_ fold change lower than *−*0.5 To assess whether positively selected genes were more frequently up-regulated or down-regulated under stress conditions, we compared the proportions of differentially expressed genes between positively selected and non-selected genes using Fisher’s exact tests for each condition.

### 2.8 Functional Annotation of candidates genes

To assess whether our candidate genes are enriched for agronomically relevant functions, we performed functional annotation analyzes, including gene ontology (GO) assignment, identification of immunity and abiotic stress-related genes, and protein–protein interaction network analysis. Because the remaining species yielded too few candidate genes for meaningful inference, these analyzes were restricted to *Ae. mutica* and *Ae. speltoides*.

#### 2.8.1 Go enrichment

GO enrichment analysis was performed using the dedicated tool implemented in TriticeaeExpDB (T. Li et al. 2026). For *Ae. speltoides*, which has a direct annotation in the database, we used its own gene identifiers. For *Ae. mutica*, which lacks direct annotation, we mapped each gene to its ortholog in *T. aestivum* and used the corresponding *T. aestivum* annotation.

#### 2.8.2 Identification of genes with known functions

For candidate genes also showing lineage-Specific acceleration in non-synonymous substitution rates, we used InterProScan (InterPro 108.0) (Blum et al. 2025) to obtain detailed protein domain. To assess the potential involvement of candidate genes in biotic and abiotic stress responses, we performed BLAST searches against two specialized databases using the following common criteria: E-value *≤* 10*^−^*^5^, identity *≥* 65%, alignment length *≥* 50; for each query we retained the best hit based on the lowest E-value.

### Immunity–related genes

We used the Pathogen Receptor Genes database (PRGdb) (Calle García et al. 2022) with the above BLAST criteria to identify pathogen receptor genes. Independently of BLAST annotation, we screened all genes with DRAGO3 (Calle García et al. 2022) to detect conserved resistance-related domains (e.g. leucine-rich repeats (LRR), kinase, NBS, TIR). Genes containing at least one such domain were classified as immunity-related. The enrichment of immunity-related genes among positively selected genes was assessed using Fisher’s exact tests.

### Abiotic stress resistance genes

To identify abiotic stress resistance genes, the PlantASRG database (H. Zhang, X. Liu, et al. 2025) was used as a reference for BLAST searches. Due to the low number of hits, no enrichment analysis could be performed.

### 2.9 Protein–protein interaction network analysis of positively selected genes

To test whether our candidate genes are more interconnected than expected by chance, we performed two complementary network analyzes using the STRING database (v12.0) (Szklarczyk et al. 2023). For each focal species, we first mapped the candidate gene identifiers to their wheat orthologs (STRING IDs).

#### 2.9.1 STRING web-based enrichment analysis

We submitted the list of our candidate genes to the STRING web interface, which computes a protein-protein interaction enrichment analysis by comparing the number of observed interactions among the candidates proteins to the number expected for a random set of proteins of the same size, drawn from the entire wheat reference network. Additionally, STRING performed functional enrichment (Gene Ontology, KEGG pathways, Pfam domains) using the hexaploid wheat genome as background.

#### 2.9.2 Permutation tests controlling for structural biases

To further assess whether the connectivity of candidate genes exceeds that of the background of all analyzable genes, we conducted a series of permutation tests. High-confidence interactions were retained by filtering for a combined STRING score *>* 0.7. The observed connectivity between candidate genes was measured as the density of the induced subgraph, i.e., the number of observed edges between candidate genes divided by the total number of possible unordered pairs. We tested whether candidate genes have a different number of interaction partners (degree) than non-candidate genes using a two-sided Wilcoxon rank-sum test. This addresses whether positively selected genes are systematically more or less connected in the global PPI network.

Additionally, we generated two distinct null distributions by permutation. The first was a degree- and length-matched permutation, in which random gene sets were sampled while preserving both the degree distribution (number of interaction partners) and the gene length (number of coding sites) of the candidate genes. This was achieved by stratifying the background gene pool into 10 degree bins and 5 length bins (determined by quantiles) and sampling the same number of genes from each bin as observed in the candidate set. This model controls for potential biases introduced by naturally more connected or longer genes. The second was a network rewiring approach, where the entire interaction network was randomised while keeping the degree sequence of all nodes fixed using the “keeping degseq” algorithm from the igraph package; each rewiring iteration performed 10 times the number of edges in the original network to ensure thorough mixing, and the candidate set was then treated as a fixed set of nodes. This tests whether the observed connectivity is an emergent property of the global network topology rather than a specific functional clustering of candidates.

For each null model, we performed 1000 permutations. Empirical p-values were calculated as the proportion of permuted densities that were equal to or greater than the observed density. A p-value *<* 0.05 was considered statistically significant.

### 2.10 Pathways analysis

#### 2.10.1 Pathways data

We downloaded gene sets from the KEGG (Kanehisa et al. 2025) and Plant reactome (Naithani et al. 2019) database. We extracted pathways separately from the KEGG and Plant Reactome databases, then merged the two sets, retaining both pathways that were shared between the databases and those that were specific to each. Pathways were directly retrieved from the data of *T. urartu* and *Ae. tauschii*, while for *Ae. mutica* and *Ae. speltoides* we used orthologs from *T. astivum* to retrieve the corresponding pathways. A set of genes represents a group of genes involved in biochemical interactions that underlie a biological process. Examples of gene sets include biosynthesis of secondary metabolites, cellular processes, or plant hormone signal transduction. We removed genes sets with less than eight genes, the remaining collection of 328 pathways were used for our analysis (110 for *Ae. speltoides*, 113 for *Ae. mutica*, 62 for *Ae. tauschii* and 43 for *T. urartu*).

#### 2.10.2 Gene Set Level Dn/Ds and Pn/Ps Ratios

To calculate the ratio *Dn/Ds* for a gene set, we summed up separately the nonsynonymous and synonymous fixed mutations found in all genes belonging to a given gene set and took their ratio as:

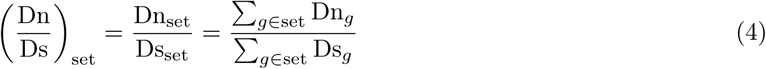

The *Pn/Ps* ratio was calculated in a similar way. We used the count obtained with the branch-specific impMKT. We then adapted the classical Fisher’s exact test for gene-level MKT to the pathway level by applying it to aggregated counts (Dn_set_, Ds_set_, Pn_set_, Ps_set_).

#### 2.10.3 Stratified Permutation Analysis

Although Fisher’s exact test identifies pathways with significant MKT deviations, the structural properties of the pathways (size, gene-length composition, and gene overlap across pathways) can affect MKT statistics independently of selection. Therefore, we performed a stratified permutation analysis, analogous to the permutation tests used for protein-protein interaction networks, to determine whether the observed pathway-level signals remain more extreme than expected after accounting for these features. The permutation was applied only to pathways that remained significant after Fisher’s test.

Each permuted pathway was constructed by sampling genes from a background pool while preserving three properties of the observed pathway: (i) the same number of genes (pathway size), (ii) the same gene-length composition (using five length strata defined from species-specific quantiles), and (iii) the same pathway-overlap structure (by weighting the sampling probability of each gene by its pathway-membership degree). For each permutation, the genes sampled were aggregated to obtain pathway-level *Pn*, *Ps*, *Dn*, and *Ds* counts, and the MK statistic was recalculated. This procedure was repeated 1000 times per pathway.

Empirical two-sided p-values were computed as:

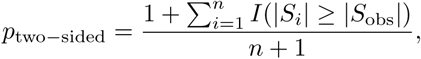

where *S_i_* is the MKT statistic from permutation *i* and *n* = 1000. This analysis tests whether the observed pathway-level MK signal remains more extreme than expected for pathways that share similar structural properties.

#### 2.10.4 Jackknife Approach to Detect Pathways with Outlier Genes

We used a jackknife approach to examine the effect of individual genes within a given gene set on our results. For each significant pathway, we repeatedly removed one gene and recalculated the *Dn/Ds* and *Pn/Ps* values. If removing a single gene caused the pathway to lose statistical significance, then such pathways were not retained as candidate gene sets for polygenic selection.

#### 2.10.5 Enrichment of candidate genes in robust pathways

Finally, to assess whether the pathway-level signals were independent of the individual candidate genes already identified, we tested for enrichment of branch-specific candidate genes among the pathways that remained significant after permutation. This allowed us to determine whether the two scales of analysis (gene-level and pathway-level) converge on the same loci or capture distinct aspects of the selection signal.

### 2.11 Use of Language Models

ChatGPT (OpenAI, GPT-5.5, accessed in 2025 through the ChatGPT web interface) was used for language editing and syntax improvement of the manuscript.

## 3 Results

### 3.1 Two species MKT

#### 3.1.1 Coverage analysis

Our analysis of transcriptomic coverage across focal species revealed that the proportion of well-covered genes was broadly comparable among the four species, ranging from 21.9% to 26.3% of the total transcriptome, with a difference of less than 5 percentage points between the highest and lowest values (Table 1). However, after applying orthology and polymorphism filters, the percentage of analyzable genes was markedly higher in outcrossing species (14.34% in *Ae. speltoides* and 10.95% in *Ae. mutica*) than in selfing species (6.63% in *Ae. tauschii* and 2.56% in *T. urartu*). This difference is consistent with the lower standing genetic variation expected under self-fertilization.

#### 3.1.2 Testing for the evidence of positive selection

Standard MKT and impMKT (cutoff 0.15) were applied to detect genes under positive selection between focal species (Table 2). Although the impMKT filtered out more genes due to its correction for SDM (resulting in fewer analyzable genes), it consistently identified a higher proportion of positively selected genes than the standard MKT. In the outcrossing species, the increase was most pronounced: in *Ae. speltoides* the proportion rose from 7.60% (MKT) to 11.58% (impMKT), and in *Ae. mutica* from 6.08% to 8.11%. In the selfing species, the absolute numbers were much lower, but the same trend was observed (e.g., *Ae. tauschii* : 0.99% vs. 1.33%).

**Table 2:**
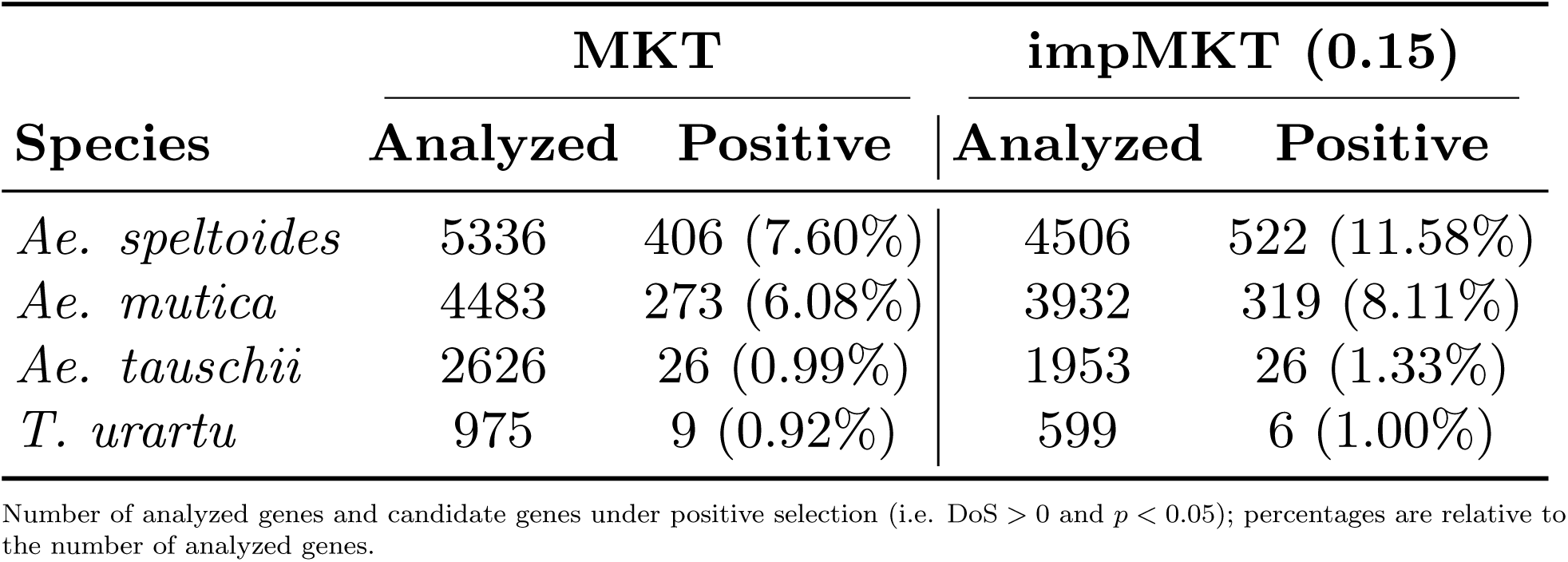
Summary of standard MKT and impMKT results across species.

| Species | MKT |  | impMKT (0.15) |  |
| --- | --- | --- | --- | --- |
|  | Analyzed | Positive | Analyzed | Positive |
| *Ae. speltoides* | 5336 | 406 (7.60%) | 4506 | 522 (11.58%) |
| *Ae. mutica* | 4483 | 273 (6.08%) | 3932 | 319 (8.11%) |
| *Ae. tauschii* | 2626 | 26 (0.99%) | 1953 | 26 (1.33%) |
| *T. urartu* | 975 | 9 (0.92%) | 599 | 6 (1.00%) |

Number of analyzed genes and candidate genes under positive selection (i.e. DoS $> 0$ and $p < 0.05$ ); percentages are relative to the number of analyzed genes.

### 3.2 Three-species MKT

#### 3.2.1 Coverage analysis

To estimate divergence on the branch of our focal species, we used CODEML with species triplets (Table 3). As expected, the triplet comprising the more closely related species *Ae. speltoides*, *Ae. mutica*, and *Ae. tauschii* yielded the highest number of orthologs (13,612), while the triplets involving more distant outgroups such as *Ae. tauschii* and *T. urartu* resulted in fewer orthologs.

**Table 3:**
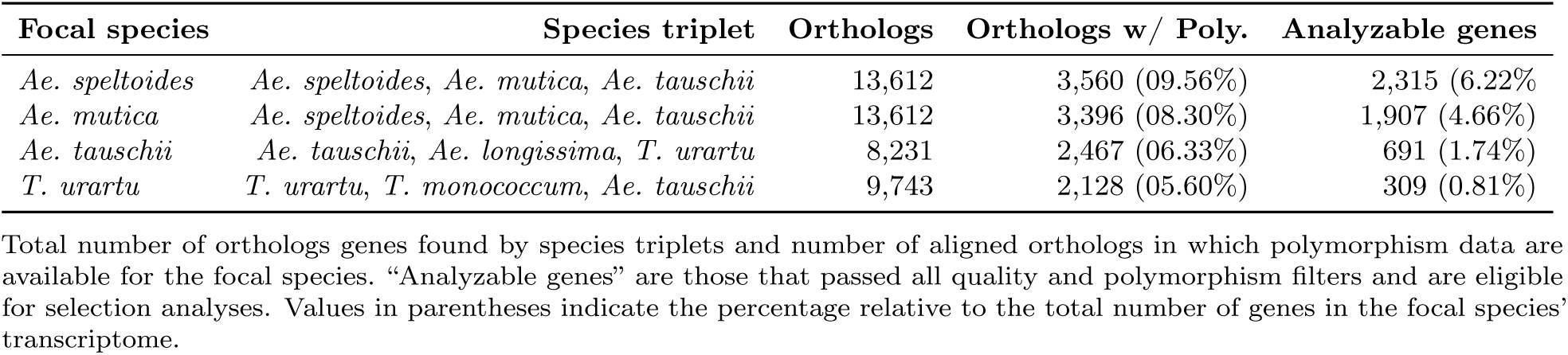
Ortholog detection and filtering statistics for the 3 species MKT analysis.

#### 3.2.2 Testing for evidence of positive selection

When restricting divergence estimates to the focal lineage, the number of analyzable genes and the proportion of positively selected candidates again differed markedly between allogamous and autogamous species (Table 4). As observed with standard approaches, *Ae. speltoides* and *Ae. mutica* showed a much higher proportion of positive candidates (1.88–5.63% of analyzable genes), while the autogamous species *Ae. tauschii* and *T. urartu* yielded almost no positively selected genes (with only two genes in *T. urartu* under branch-specific impMKT). Among allogamous species, the branch-specific impMKT (cutoff 0.15) consistently identified a larger fraction of positively selected genes than branch-specific MKT (e.g., *Ae. speltoides*: 5.63% vs. 3.15%; *Ae. mutica*: 3.14% vs. 1.88%).

**Table 4:** Summary of branch-specific MKT and branch-specific impMKT (cutoff 0.15) results across species.

| Species | Branch-specific MKT |  | Branch-specific impMKT (0.15) |  |
| --- | --- | --- | --- | --- |
|  | Analyzed | Positive | Analyzed | Positive |
| *Ae. speltoides* | 2,315 | 73 (3.15%) | 1,864 | 105 (5.63%) |
| *Ae. mutica* | 1,907 | 36 (1.88%) | 1,651 | 52 (3.14%) |
| *Ae. tauschii* | 691 | 0 (0%) | 530 | 0 (0%) |
| *T. urartu* | 309 | 0 (0%) | 204 | 2 (0.98%) |

Number of analyzed genes and candidate genes under positive selection (i.e. DoS > 0 and $p < 0.05$ ); percentages are relative to the number of analyzed genes.

Compared to the standard two-species framework (Table 2), the branch-specific approach substantially reduced the number of analyzable genes, as expected because a second outgroup is needed. Consequently absolute numbers of positive candidates were also lower. However, the relative differences between allogamous and autogamous species remained very similar, and the impMKT version still outperformed its MKT counterpart. For example, in *Ae. speltoides* the proportion of positive genes increased from 3.15% (branch-specific MKT) to 5.63% (branch-specific impMKT), a relative gain comparable to that seen with standard methods (7.6% to 11.6%).

To document the overlap and specificity of each method, we generated UpSet plots displaying the intersection sizes of positively selected genes (DoS *>* 0, *p*-value *<* 0.05) in the four MKT-based methods for each species (Supplementary Figure **??**). The UpSet plot comparisons reveal a limited concordance among MKT-based methods, indicating that signals of positive selection are highly method-dependent between species. In both *Ae. speltoides* and *Ae. mutica*, a large number of candidate genes were identified by a single method, with only a fraction shared between approaches. In particular, standard MKT and impMKT approaches tend to recover more similar sets of genes, whereas branch-specific methods identify largely non-overlapping candidates.

To further evaluate the consistency of selection signals across lineages, we restricted the analysis to orthologous genes that were analyzable by both the 2 species impMTK and the branch-specific impMKT approaches in two focal species (*Ae. mutica* and *Ae. speltoides*). We then generated an UpSet plot summarizing the overlap of significant genes across methods and species (Figure 3). The comparison revealed marked differences between the two approaches. Among the 53 genes identified by the impMKT in *Ae. mutica*, 19 were also detected in *Ae. speltoides* (35.8%), suggesting a substantial proportion of apparently convergent adaptive signals. Similarly, 19 of the 28 significant genes detected in *Ae. speltoides* were shared with *Ae. mutica* (67.9%). In contrast, the branch-specific analysis identified only two shared genes among the 18 significant genes detected in each species (11.1%). These differences are expected because the two methods rely on different evolutionary contrasts. In the impMKT, each species is compared against the same set of divergence events, while only the polymorphism data differ between species. Consequently, adaptive signals originating from substitutions that occurred before lineage divergence may be detected independently in multiple species, creating an apparent pattern of convergence. In contrast, the branch-specific impMKT uses branch-specific divergence estimates and explicitly assigns substitutions to individual evolutionary lineages. As a result, adaptive signals can be attributed to a single species, leading to a much lower number of shared candidate genes. The results suggest that the relatively high level of overlap observed with the impMKT may partly reflect limitations in the ability of this framework to localize adaptive substitutions to particular lineages. The branch-specific approach provides a more conservative estimate of convergent adaptation by distinguishing lineage-specific from shared historical divergence events. However, the number of orthologous gene pairs that could be analyzed by both methods was limited, and thus the observed patterns should be interpreted with caution.

**Figure 3:**
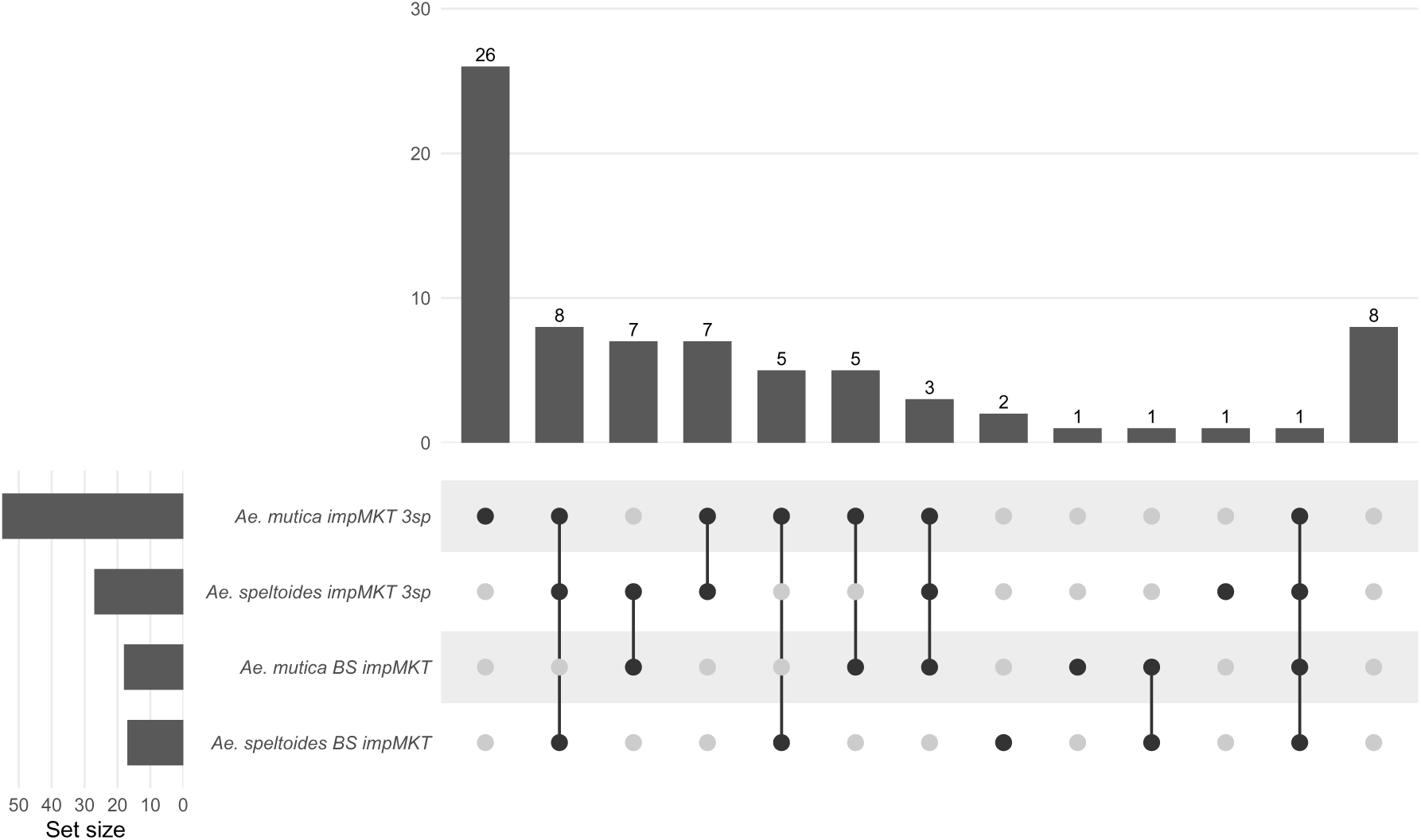
Comparison of significant genes detected by MKT-based methods across the two allogamous focal species *Ae. speltoides* and *Ae. mutica*. UpSet plots display the intersection sizes of genes with DoS *>* 0 and p-value *<* 0.05 for each method combination. The analysis includes all genes that were analyzed by both methods on both species (total of 75 genes). The horizontal bar plots (left) show the total number of significant genes per method, while the vertical bar plots (top) indicate the number of genes shared between method intersections. Each column represents a specific combination of methods, with connected dots indicating which methods are included in the intersection. Columns containing only grey dots correspond to genes that were not candidate in any method (i.e., not significantly detected by any of the four approaches). MKT 2sp = two species MKT, impMKT 2sp = two species impMKT, BS MKT = branch-specific MKT, BS impMKT = branch-specific impMKT.

### 3.4 Association Between adaptative evolution and Signatures of Accelerated Protein Evolution

Branch-specific MKT/impMKT combines polymorphism and divergence data to infer lineage-specific adaptive evolution. Importantly, both branch-specific MKT/impMKT and lineage-specific *dN/dS* analyzes rely on the same underlying divergence signal (nonsynonymous and synonymous substitutions), but branch-specific MKT/impMKT additionally incorporates within-species polymorphism as a neutral reference framework. This allows divergence to be interpreted relative to the level of polymorphism expected under neutrality along each lineage. The branch specific version of the MKT is designed to identify genes showing an excess of nonsynonymous divergence relative to neutral expectations. Consequently, if divergence alone captures most of the adaptive signal, our candidates should overlap with genes exhibiting elevated *dN/dS* ratios or signatures of lineage-specific acceleration detected from divergence-based analyzes. In that case, most signature of adaptive evolution could potentially be inferred without polymorphism data and would allow to analyses genes with little or no polymorphism, for example due to selective sweeps, hitchhiking effects, or limited sequence coverage. Conversely, limited overlap would indicate that polymorphism provides additional information that cannot be recovered from divergence alone. To evaluate this hypothesis, we applied CODEML branch-model likelihood-ratio tests to identify genes with significant acceleration on the focal branch. In *Ae. speltoides*, 86 genes were classified as accelerated, and 21 (24.4%) of them were BSimpMKT candidates (Fisher’s exact test: OR = 6.23, *p* = 3.3 *×* 10*^−^*^9^). In *Ae. mutica*, 68 genes were classified as accelerated, and 16 (25.8%) were BSimpMKT candidates (OR = 12.04, *p* = 7.8 *×* 10*^−^*^11^). Despite these significant enrichments, the overlaps remained limited: most BSimpMKT candidates were not captured by divergence-based acceleration alone. Thus, polymorphism data provide essential information for detecting adaptive evolution that cannot be recovered from divergence-based methods alone.

### 3.5 Expression Analysis: Abiotic and Biotic Stress of Candidate Genes Under Positive Selection

Across multiple stress conditions, we compared the proportion of up-regulated genes between our candidates and background genes. In *Ae. mutica*, a significant enrichment of up-regulated genes was observed under *Zymoseptoria* infection at 9 days post-inoculation (Fisher’s exact test, *p <* 0.01; (Supplementary Figure **??**). No significant differences were detected at earlier or later time points. In *Ae. speltoides*, candidate genes showed a significantly higher proportion of up-regulated genes under powdery mildew infection, particularly at 24 and 72 hours post-inoculation (*p <* 0.05 and *p <* 0.01, respectively; Supplementary Figure **??**). In contrast, no significant enrichment was observed under strip rust infection. Although we observed significant enrichment under fusarium stress at 36 hours and 48 hours. Across other stress conditions tested, no consistent or significant differences in expression patterns were detected between candidate and non-candidate genes, neither for up-regulated nor down-regulated genes.

### 3.5 Functional Annotation

#### 3.5.1 Go enrichment

After correction for multiple tests, no Gene Ontology (GO) term reached statistical significance (adjusted p-value < 0.05) in any species when considering candidate genes. In *Ae. speltoides*, the lowest adjusted p-value was 0.068, associated with positive reaction to stimulus.

#### 3.5.2 Functional characterization of candidate genes

We investigated the functional annotation using InterProScan of the 21 genes of *Ae. speltoides* and 15 genes of *Ae. mutica* that both showed signatures of accelerated protein evolution and were classified as candidate genes. The *Ae. speltoides* candidate *EVM0020846.1* encodes a predicted protein kinase that contains an interleukin-1 receptor-associated kinase-like domain STKc_IRAK. Furthermore, the data base prosite detected a serine/threonine protein kinases active-site signature, and a protein kinases ATP-binding region. Interestingly, a wheat stem rust resistance gene *Sr43* (Yu et al. 2023) and resistance genes against *Septoria tritici* blotch *Stb16q* (Saintenac et al. 2021) both contain STKc_IRAK kinase domains, supporting the hypothesis that *EVM0020846.1* may participate in an immune or stress-related signaling response. An additional annotation through CATH associated this protein with FERONIA-like receptor kinases, a family known to regulate plant immunity, cell wall integrity, mechanosensing, and abiotic stress responses (Cheung 2024). Another *Ae. speltoides* candidate, *EVM0020069.1*, contains LRR, which are commonly associated with extracellular recognition and immune signaling in plants (Klymiuk et al. 2021). Among the remaining candidates, two genes are particularly notable. *EVM0049693.1* contains an alcohol dehydrogenase (ADH) domain, a gene family implicated in waterlogging tolerance during wheat seed germination (Shen et al. 2021). *EVM0052755.2* encodes a Glycoside Hydrolase Family 1 (GH1) -glucosidase, a class of enzymes involved in plant defense metabolism and broadly induced under abiotic stresses such as drought and cold (Alwutayd et al. 2026). In *Ae. mutica*, candidate gene *Ammut_EIv1.0_0092140.1* also encodes a kinase containing an STKc_IRAK domain, together with signatures characteristic of active serine/threonine kinases. Additional annotations identified similarity to MAP3K-like kinases, components of conserved MAP kinase signaling cascades that regulate plant growth, development, and responses to biotic and abiotic stresses (Huang et al. 2025; Shu et al. 2025). The *Ae. mutica* candidate *Ammut_EIv1.0_0407820.1*, contains SEC14 and CRAL/TRIO lipid-binding domains. SEC14-like proteins are involved in membrane trafficking and phospholipid-mediated signaling, and members of this family in wheat have been implicated in salt stress tolerance (Qin et al. 2023). The candidate *Ammut_EIv1.0_0158190.1* contains a cytochrome P450 domain. cytochrome P450s play a role in the host response to diseases, including the wheat response to Fusarium head blight (FHB) disease (Walter and Doohan 2011). In wheat, a cytochrome P450 gene of the CYP72A subfamily (*TaCYP72A*) is induced in response to Fusarium mycotoxin deoxynivalenol (DON), and gene-silencing experiments have shown that it contributes to DON resistance and grain development (Gunupuru et al. 2018). Although *Ammut_EIv1.0_0158190.1* cannot be directly linked to these functions based solely on domain annotation, the involvement of P450 proteins in pathogen responses and stress-related processes makes this positively selected gene an interesting candidate for future functional characterization. For the two candidates of *T. urartu* one of them *TuG1812G0300002508.01.T01* contains a WD40 repeat domain, a motif found in stress-associated proteins such as TaWD40D in wheat (Kong et al. 2015).

#### 3.5.3 Identification of immunity-related genes and abiotic stress resistance gene

To assess the functional relevance of our candidate genes, we compared their protein sequences with two curated databases: PRGdb (Supplementary Tables **??** and **??**) that catalog plant resistance genes, and PlantASRG (Tables 5), which focus on genes involved in abiotic stress tolerance. For each candidate, we retained the best BLAST hit and report its sequence identity and functional annotation when available. Overall, we retrieved 28 best BLAST hits for *Ae. speltoides* and 19 for *Ae. mutica* from the PRGdb database. We also assessed if the candidate genes exhibited a higher proportion of immunity-related domains compared to non-selected genes in both species. In *Ae. speltoides*, 9.92% of positively selected genes carried immunity-related domains versus 5.26% of non-selected genes (Fisher’s exact test, *p* = 0.02996). In *Ae. mutica*, a similar trend was observed (12.90% vs 5.65%), (Fisher’s exact test, *p* = 0.0.0262).

**Table 5:** Best BLAST hits of *Ae. mutica* and *Ae. speltoides* candidate genes against the PlantASRG database.

| Gene ID | Gene name | Species | Identity (%) | Stress type |
| --- | --- | --- | --- | --- |
| *Ae. speltoides* |  |  |  |  |
| EVM0010149.1 | TaMYB3R1 | *Triticum aestivum* | 97.686 | Drought |
| EVM0042405.1 | Ta-Ub2 | *Triticum aestivum* | 97.368 | Drought |
| EVM0039720.1 | EaGly III | *Erianthus arundinaceus* | 85.530 | Drought |
| EVM0022205.1 | OsTMF | *Oryza sativa* | 78.215 | Cold |
| *Ae. mutica* |  |  |  |  |
| Ammut_EIv1.0_0634380.1 | BAM1 | *Arabidopsis thaliana* | 70.684 | Drought |
| Ammut_EIv1.0_0205290.1 | MSZEP | *Medicago sativa* | 68.489 | Drought |
| Ammut_EIv1.0_0905850.1 | OsMAR1 | *Oryza sativa* | 66.271 | Salt and alkali |
| Ammut_EIv1.0_0788330.1 | StAPX | *Solanum lycopersicum* | 65.526 | Salt and alkali |

For each gene, PlantASRG matches include the gene name, source species, identity percentage, and stress type.

PlantASRG analyzes identified a small subset of candidate genes associated with abiotic stress responses. In *Ae. speltoides*, four genes were linked to drought and cold stress, including *TaMYB3R1* (97.7% identity) and *OsTMF* (78.2%). In *Ae. mutica*, four genes were also identified, associated with drought (e.g., *MSZEP*, *BAM1* ) and salt/alkali stress (e.g., *StAPX*, *OsMAR1* ). These results highlight the involvement of positively selected genes in stress-related functions.

We investigated the functional annotation of genes that both showed signatures of accelerated protein evolution and were classified as candidate genes. 5 of 21 such genes in *Ae. speltoides* and 7 of 15 in *Ae. mutica* matched entries in the Pathogen Receptor Genes database. And 1 gene of *Ae. speltoides* and 1 of *Ae. Mutica* in PlantASRG database.

#### 3.5.4 Protein–Protein Interaction Network Analysis

We assessed whether our candidate genes tend to form a more interconnected protein–protein interaction (PPI) network than expected by chance, using two complementary approaches. Candidate STRING IDs were submitted to the STRING web interface (v12.0), to assess protein–protein interaction (PPI) enrichment and functional enrichment. Neither species showed a significant excess: in *Ae. mutica* (53 proteins, 33 observed vs. 29 expected interactions, PPI enrichment *p* = 0.27); in *Ae. speltoides* (113 proteins, 115 observed vs. 114 expected, *p* = 0.489). Thus, candidate sets are not globally more interconnected than random gene sets of the same size. The most significant functional enrichments in *Ae. mutica* were related to cellular anatomical, DNA repair–related regulatory pathways, suggesting a possible enrichment of functions involved in genome maintenance (see supplementary file File_S4_GO_enrichment_mutica.tsv in the Zenodo repository at https://doi.org/10.5281/zenodo.21397724. In *Ae. speltoides* several functional terms were significantly over-represented. Among these, the most notable were reactome pathways related to infectious disease (MAP-5663205, 5 of 153 proteins, *p* = 7.8 *×* 10*^−^*^4^) and disease (MAP-1643685, 5 of 190 proteins, *p* = 0.0011); see supplementary file File_S3_GO_enrichment_speltoides.tsv in the Zenodo repository at https://doi.org/10.5281/zenodo.21397724). Because the STRING PPI enrichment test does not control for gene-level properties (degree, length) or global network topology, we performed additional permutation tests on the high-confidence STRING subnetwork (combined score *>* 0.7). We performed two permutation tests. The degree- and length-matched permutation (stratifying background genes into 10 degree bins and 5 length bins) showed that candidate genes in both species are more interconnected than expected for random genes with similar degree and length (*Ae. speltoides*: *p* = 0.01; *Ae. mutica*: *p <* 0.005). The network rewiring test (randomising edges while preserving node degrees) tests whether the observed connectivity reflects global network topology rather than specific functional clustering. After rewiring, enrichment remained significant only in *Ae. mutica* (*p* = 0.025), but not in *Ae. speltoides* (*p* = 0.355). Taken together, these results suggest that while candidate genes in both species tend to be more connected than expected from their individual degree and length, only in *Ae. mutica* does this connectivity exceed expectations based on the global network topology. Hence, *Ae. mutica* may harbor a more robust subnetwork of interaction genes.

Several individual interacting candidate genes stood out due to their known biological functions and specific interaction partners. In *Ae. speltoides*, *ZFP-1* encodes a zinc finger protein that plays an important role in plant growth, development, and biotic and abiotic stress responses (B. Sun et al. 2019; A. Sun et al. 2022). It interacts with a ubiquitin-like domain-containing protein – itself a candidate gene. Ubiquitination is a major post-translational modification involved in diverse processes including photomorphogenesis, vascular differentiation, flower development, phytohormone and light signalling, and stress responses (W. Gao et al. 2024). Another candidate gene identified in the network is *ATG16b*, autophagy-related proteins (ATGs) are an indispensable biological process and play crucial roles in plant growth and plant responses to biotic and abiotic stresses (Yue et al. 2022). *ATG16b* also interacts with other candidate genes, including protein kinase domain-containing protein these proteins with a tandem kinase structure have recently emerged as immune resistance in cereal crops (Bernasconi et al. 2025). In *Ae. mutica*, the set of interacting positively selected genes also includes several proteins with relevant functional annotations. Zeaxanthin epoxidase (ZE), a chloroplastic enzyme involved in the xanthophyll cycle and ABA-mediated stress responses, was found to interact with other candidate genes. In wheat, suppression of ZEAXANTHIN EPOXIDASE 1 increases H_2_O_2_ accumulation and restricts stripe rust growth, directly linking ZE to disease resistance (Chang et al. 2023). More generally, ZE plays multifaceted roles in photoprotection, ABA signalling, and carotenoid metabolism (Chang et al. 2023). In the network, ZE interacts with another candidate gene a terpene cyclase/mutase family member – a class of enzymes that may produce phytoalexins or defence-related signalling molecules (Polturak et al. 2022).

### 3.6 Pathways analysis

We tested 113 pathways in *Ae. speltoides*, 110 in *Ae. mutica*, 62 in *Ae. tauschii*, and 43 in *T. urartu*. Two-sided Fisher’s exact tests identified 62 pathways in *Ae. speltoides* and 21 in *Ae. mutica* showing an excess of nonsynonymous divergence (*Dn/Ds > Pn/Ps*, *p <* 0.05; see Supplementary Tables S1 and S2 in the supplementary PDF). No significant pathways were detected in *Ae. tauschii* or *T. urartu*. Pathways related to metabolism and genetic information processing were frequently observed among the significant pathways. Some examples of defense related pathways were still detected as Responses to stimuli: abiotic stimuli and stresses and Plant-pathogen interaction in *Ae. speltoides*. After applying the stratified permutation analysis to control for the structural properties of the pathway (pathway size, gene length composition - and structure of the overlap of the pathway) to the pathways that were significant in the Fisher test, and after evaluating the robustness to single-gene effects using jackknife analyzes, only a subset remained significant: 11 in *Ae. speltoides* and 1 in *Ae. mutica* (Table 7).

Among these pathways, several are related to biological functions likely to be important for environmental adaptation and cellular maintenance in Triticeae species. In *Ae. speltoides*, the pentose phosphate pathway plays a protective role against low-temperature stress during wheat reproduction (Dai et al. 2026). Similarly, the protein processing in the endoplasmic reticulum pathway plays a central role in mitigating cold-induced protein misfolding and cellular stress responses (A.-m. Zhang et al. 2023). The nucleotide excision repair pathway, another candidate supported, is responsible for the removal of UV-induced DNA lesions and is therefore critical for maintaining genome integrity under elevated exposure to solar radiation (Sancar 2016; Oztas et al. 2018). Interestingly, the protein export pathway was identified in both *Ae. speltoides* and *Ae. mutica*. This pathway is essential for the transport of storage proteins into the developing endosperm and directly influences grain protein accumulation and grain quality traits (Hasrak et al. 2026).

To test whether these pathway-level signals reflected the same loci already identified as our candidate genes, we assessed the enrichment of candidate genes among the members of pathways that passed all four layers of evidence. In *Ae. speltoides*, 12 of 40 annotated candidates (30 %) fell within one of the 11 robust pathways, compared to 18.4 % expected by chance (OR = 1.96; one-sided Fisher’s exact p = 0.048, two-sided p = 0.062). This enrichment was not detected at the Fisher-only threshold (69 pathways; OR = 2.21, p = 0.128), confirming its specificity for the validated subset. However, five of the 11 robust pathways did not contain an individually significant candidate gene, indicating that part of the pathway-level signal is not captured by individual gene signals. In *Ae. mutica*, the single robust pathway (Protein export) contained no branch-specific candidate gene, though only one robust pathway was recovered in this species, preventing a comparably powered test.

## 4 Discussion

### 4.1 Evidence of positive selection in WWR

We detected marked differences among species in the proportion of genes inferred to be under positive selection, with consistently higher values in the outcrossing species (*Ae. speltoides* and *Ae. mutica*) than in the predominantly selfing species (*Ae. tauschii* and *T. urartu*). Continually, branch-specific MKT and impMKT identified substantially more candidate genes in the two outcrossing species (up to 3.15% and 5.63% in *Ae. speltoides*, and 1.88% and 3.14% in *Ae. mutica*) than in the selfing species, where almost no candidates were detected. A similar pattern was seen at the pathway level, with significant positive selection signature only found in out-crossing species.

These differences are expected given the strong effect of mating system on the efficacy of selection. The selective strength acting on mutations is proportional to *N_e_s*, where *N_e_* is the effective population size and *s* the selective advantage of a beneficial mutation (Ewens 2004). In complete self-fertilization, *N_e_* is reduced by at least half, weakening the efficacy of selection (Nordborg 1997), because newly arising beneficial mutations are more susceptible to stochastic loss through genetic drift in selfing populations (Glémin 2012; Hartfield et al. 2017). Selfing also amplifies the effects of linked selection genome-wide through reduced effective recombination (Burgarella et al. 2024). Increased linkage among sites enhances Hill–Robertson interference, thereby reducing the efficacy of selection against deleterious mutations and limiting the fixation probability of beneficial alleles. Reduced effective population size may additionally constrain adaptation from standing genetic variation by limiting the pool of segregating variants from which beneficial alleles can arise (Glémin and Ronfort 2013; Hartfield et al. 2017).

The patterns observed here in WWR are in line with Burgarella et al. (2024) which did not detect positive selection signatures in *T. urartu* and *Ae. tauschii*. Similarly in other plant systems outcrossing species *Capsella grandiflora*, approximately 40% of amino-acid substitutions were estimated to be fixed by positive selection, together with more efficient purging of deleterious mutations (Slotte et al. 2010; Williamson et al. 2014). By contrast, the predominantly selfing *Arabidopsis thaliana* showed little evidence of adaptive fixations and instead accumulated SDMs (Kim et al. 2007).

Overall, both gene-level and pathway-level analyzes support the view that the mating system strongly shapes the efficacy of selection in WWR, with outcrossing lineages exhibiting stronger signatures of positive selection and selfing lineages showing patterns consistent with the accumulation of SDMs.

### 4.2 Effect of SDM

Our results highlight the trade-off between accounting for SDMs and retaining sufficient polymorphism for MKT inference. By incorporating SDMs, impMKT reduces potential bias, but also decreases the number of analyzable genes, particularly in species with low polymorphism, as previously reported by Murga-Moreno, Coronado-Zamora, Casillas, et al. (2022). However, this filtering increased the number of candidate genes detected in outcrossing species, whereas no benefits were observed in predominantly selfing species. These results suggest that impMKT is most informative in species with moderate to high polymorphism, while more conservative thresholds may further reduce statistical power in selfing species. The 15% allele frequency threshold used here, following Charlesworth and Eyre-Walker (2008), may therefore require adjustment, and future simulation-based studies could help define species-specific thresholds that better balance SDM exclusion and polymorphism retention.

### 4.3 Effect of branch-specific divergence estimation

While the standard two-species MKT versions allow a larger number of genes to be analyzed, divergence is estimated from pairwise differences between the focal species and an outgroup. Consequently, nonsynonymous and synonymous substitutions accumulated along both lineages since their common ancestor are pooled into the same *Dn* and *Ds* counts. Adaptive signals inferred by the test therefore cannot be unambiguously assigned to the focal lineage and may partly reflect substitutions that occurred in the outgroup. To overcome this limitation, we implemented a branch-specific framework using three species and the codeml program from the PAML package (Z. Yang 2007). This approach estimates *dN* and *dS* directly along the focal branch under a phylogenetic maximum-likelihood model, allowing substitutions to be assigned to individual lineages. As a result, adaptive evolution can be localized to specific branches rather than inferred from divergence accumulated across multiple lineages. In addition, this framework accounts for multiple substitutions at the same site (Z. Yang 2007), providing more accurate branch-specific estimates of divergence. This difference had a substantial impact on the inferred patterns of adaptation. In the standard impMKT analysis, a large proportion of candidate genes appeared to be shared between *Ae. mutica* and *Ae. speltoides*, on the contrary, branch-specific impMKT identified only two shared genes among the 18 significant genes detected in each species (11.1%). These results indicate that many apparently convergent signals identified by the standard impMKT likely reflect substitutions that cannot be assigned to a particular lineage rather than independent adaptive evolution in both species.

Overall, the branch-specific framework provides a more conservative and biologically interpretable estimate of lineage-specific adaptation by explicitly assigning substitutions to evolutionary lineages. This increased lineage resolution comes at the cost of a reduced number of analyzable genes, owing to the requirement for high-quality multispecies orthologous alignments.

### 4.4 Gene-level vs pathway level detection

Although the MKT framework provides a powerful approach for identifying candidate genes under positive selection, its gene-level application has inherent limitations that may reduce the detection of adaptive evolution. First, selective pressures can vary between exons or functional domains within the same gene. For example, a protein may contain rapidly evolving regions involved in pathogen recognition together with highly conserved structural or catalytic domains. By aggregating polymorphism and divergence across the entire coding sequence, gene-level MK analyzes may, therefore, dilute localized adaptive signals. Exon-level or sliding-window approaches could help identify more specific signatures of adaptation.

A second limitation is the dependence of MKT inference on the availability of polymorphism data. Reduced coverage, limited sample sizes, or low diversity can decrease statistical power, as observed in selfing species with fewer analyzable genes. Although MKT-based approaches remain preferable when sufficient polymorphism is available because they incorporate segregating SDMs, divergence-based methods such as PAML (Z. Yang 2007) can provide complementary information when polymorphism is limited. Although they cannot directly distinguish adaptive evolution from relaxed purifying selection or other sources of rate variation (Moutinho et al. 2020; Zwonitzer et al. 2023), the overlap between accelerated-evolution genes and MKT candidates suggests that divergence-based signals can still help prioritize adaptive candidates.

At the same time, selection can act not only on individual genes, but also on groups of functionally related genes. Consequently, studies of adaptive evolution have increasingly shifted from single-gene approaches to pathway- and network-level analyzes, reflecting the recognition that many adaptive traits have a polygenic architecture and emerge from interactions between multiple genes rather than isolated loci (Daub et al. 2015; Boyle et al. 2017; Nguyen et al. 2018; Barghi et al. 2020). This shift is biologically motivated because selection acting on complex traits is expected to affect multiple components of a pathway or network. Moreover, adaptive evolution may involve many mutations of small individual effects, generating weak signals at individual loci but detectable cumulative signals across functional modules. Our gene-set MKT analysis in *Ae. speltoides* illustrates this expectation: gene-level candidates were not significantly enriched in robust pathways, and 5 of the 11 retained pathways contained no candidates, suggesting that the two approaches capture distinct aspects of selection signals. However, pathway-level analyzes also have inherent limitations. Although grouping genes increases statistical power by allowing inclusion of genes with undefined *Dn/Ds* or *Pn/Ps* ratios, it can also combine genes with different recombination contexts, GC content, and gene densities. Because evolutionary forces can act differently among genes within the same pathway, this aggregation may dilute biological signals.

Consequently, approaches at the gene and pathway-levels should be viewed as complementary rather than competing strategies. Gene-level analyzes are well suited for identifying strong-effect adaptive candidates and mechanistically interpretable loci, whereas pathway-level analyzes may better capture diffuse polygenic adaptation distributed across multiple interacting genes. Combining both scales of analysis may, therefore, provide a more complete picture of adaptive evolution.

### 4.5 Gene function

To assess the functional relevance of positively selected genes identified in *Ae. speltoides* and *Ae. mutica*, we combined GO enrichment analyzes, protein domain annotation, protein–protein interaction networks, homology searches against curated resistance databases, and pathway-level analyzes. This multi-layered approach allowed us to assess whether the selection signals detected at different scales converge on known stress-response functions, and whether they point to genes already characterized in wheat or to potentially novel candidates.

Immunity-related domains, including NB-ARC, LRRs, and kinase domains, were more frequent among positively selected genes than among non-selected genes (11.4% vs 6.4% in *Ae. speltoides*; 15.4% vs 6.9% in *Ae. mutica*), supporting a non-random association between positive selection and immunity-related functions.

Homology searches against PRGdb and PlantASRG identified candidates related to resistance and stress-associated genes. Some showed a high similarity to wheat genes, including *TaMYB3R1* and *Ta-Ub2* in *Ae. speltoides* (>97% amino acid identity), potentially representing adaptive alleles at loci already present in wheat, although causal polymorphisms need to be identified. Others genes were more similar to stress-related genes characterized in other plant species, including homologs of *BAM1*, *OsMAR1*, *StAPX*, and *MSZEP*, representing potential novel candidates in WWR. Several candidates also contained stress- and immunity-associated domains, including STKc_IRAK kinases, SEC14 lipid-binding domains, and LRRs. The STKc_IRAK domain was identified in a candidate for both *Ae. speltoides* and *Ae. mutica* and is also present in wheat resistance genes such as *Sr43* and *Stb16q* (Y. Li et al. 2025). Although domain presence alone does not demonstrate resistance function, these candidates represent priorities for future validation.

Beyond individual candidates, network and pathway analyzes revealed enrichment of immune- and stress-response functions, including disease-response and "Plant–pathogen interaction" pathways in *Ae. speltoides* and the "MAPK signaling pathway-plant" in *Ae. tauschii*, suggesting that positive selection has acted on broader stress-response networks.

Overall, our results support MKT-based approaches as a first-pass framework to prioritize candidate adaptive genes in CWRs, consistent with previous studies reporting enrichment of immune-related functions among positively selected genes in plants (Roth and Liberles 2006; Rech et al. 2012; Poppe et al. 2015). Our shortlist of priority candidates (Table 6) includes homologs of known wheat resistance genes and genes representing potentially unexplored adaptive variation in WWR, providing targets for further characterization from a crop improvement perspective.

**Table 6:** Short list of priority genes for further characterisation in *Ae. speltoides* and *Ae. mutica*. For each gene, available protein-domain annotations and orthologs in *T. aestivum* are indicated.

| Gene ID | Protein domain | *T. aestivum* ortholog |
| --- | --- | --- |
| *Ae. speltoides* |  |  |
| EVM0007799.1 | ABC1 atypical kinase-like domain | TraesCS4B02G035800.1 |
| EVM0012244.1 | AR_FR_like_1_SDR_e | TraesCS4B02G301500.1 |
| EVM0018390.1 | Myb-type | TraesCS3B02G367500.1 |
| EVM0020069.1 | At1g61320/AtMIF1, LRR domain | TraesCS5B02G280500.2 |
| EVM0020846.1 | STKc_IRAK | TraesCS2B02G484700.1 |
| EVM0022480.1 | DEAD-box | TraesCS6B02G262800.1 |
| EVM0024854.1 | TM9SF | TraesCS6B02G393800.1 |
| EVM0026538.1 | RPA1 | TraesCS4D02G246300.1 |
| EVM0027825.1 | LRR | TraesCS2A02G360300.1 |
| EVM0028151.1 | Sec23/Sec24 | TraesCS4B02G085800.1 |
| EVM0039207.1 | ABC transporter | TraesCS2B02G485600.1 |
| EVM0039894.1 | Autophagy protein 16 | TraesCS5D02G394200.1 |
| EVM0042405.1 | Ubiquitin | TraesCS1B02G241200.1 |
| EVM0045214.1 | Vaccinia Virus protein VP39 | TraesCS3B02G072500.1 |
| EVM0046547.1 | Ypt/Rab-GAP | TraesCS5B02G301700.1 |
| EVM0049109.1 | NAC | TraesCS1B02G248000.1 |
| EVM0049693.1 | Alcohol dehydrogenase (ADH) | TraesCS5B02G202800.1 |
| EVM0049852.1 | Clathrin heavy-chain (CHCR) | TraesCS4B02G049500.1 |
| EVM0052583.1 | DUF565 | TraesCS2B02G365700.1 |
| EVM0052755.2 | β-Glucosidase (GH1) | TraesCS3B02G430000.2 |
| EVM0057480.2 | ArfGap | TraesCS4A02G323700.1 |
| *Ae. mutica* |  |  |
| Ammut_EIv1.0_0029920.1 | Membrane-bound O-acyltransferase (MBOAT) | TraesCS1A02G086900.1 |
| Ammut_EIv1.0_0092140.1 | Serine/threonine-protein kinase, STKc_IRAK | TraesCS1B02G351900.1 |
| Ammut_EIv1.0_0149350.1 | DNA-directed RNA polymerases I, II, and III subunit RPABC3 | TraesCS2D02G111300.1 |
| Ammut_EIv1.0_0158190.1 | Cytochrome P450 | TraesCS2B02G167500.1 |
| Ammut_EIv1.0_0172170.1 | LRR | TraesCS2D02G207200.1 |
| Ammut_EIv1.0_0205290.1 | Forkhead-associated (FHA) | TraesCS2D02G314900.1 |
| Ammut_EIv1.0_0266380.1 | Transcription factor TFIIH subunit p52/Tfb2 | TraesCS2D02G599500.1 |
| Ammut_EIv1.0_0286740.1 | PB1 domain | TraesCS3A02G051800.1 |
| Ammut_EIv1.0_0407820.1 | CRAL/TRIO domain | TraesCS3B02G538200.1 |
| Ammut_EIv1.0_0632870.1 | Alpha-N-acetylglucosaminidase | TraesCS5B02G405200.2 |
| Ammut_EIv1.0_0652780.1 | Serine/threonine-protein kinase BSK | TraesCS5D02G501100.1 |
| Ammut_EIv1.0_0786480.1 | MORN (Membrane Occupation and Recognition Nexus) repeat | TraesCS6A02G406600.3 |
| Ammut_EIv1.0_0856250.1 | Squalene cyclase, Terpenoid cyclases/protein prenyl-transferase alpha-alpha toroid | TraesCS7D02G233100.2 |
| Ammut_EIv1.0_0880010.1 | Serine/threonine-protein kinase | TraesCS7B02G220500.1 |
| Ammut_EIv1.0_0910000.1 | DUF5600 | TraesCS7D02G430800.1 |
| *T. urartu* |  |  |
| TuG1812G0300002508.01.T01 | WD40 | TraesCS3A02G201200.1 |
| TuG1812G0700000332.01.T01 | Vaccinia Virus protein VP39 | TraesCS7A02G031100.1 |

Each candidate was identified by branch-specific MKT/impMKT analyses and shows signatures of accelerated protein evolution.

**Table 7:** Candidate Pathways after stratified permutation analysis and evaluating robustness to single-gene effects using jackknife analyses in *Ae. speltoides* and *Ae. mutica*.

| Species | Gene Set | Size (Genes) | Length (kb) | Dn/Ds | Pn/Ps | *P* | *q* |
| --- | --- | --- | --- | --- | --- | --- | --- |
| *Ae. speltooides* | Protein processing in endoplasmic reticulum | 46 | 71,04 | 0,31 | 0,14 | $1,00 \times 10^{-9}$ | $3,17 \times 10^{-8}$ |
| *Ae. speltooides* | Nucleotide excision repair | 17 | 24,96 | 0,23 | 0,06 | $3,65 \times 10^{-9}$ | $7,82 \times 10^{-8}$ |
| *Ae. speltooides* | Basal transcription factors | 11 | 13,37 | 0,55 | 0,17 | $2,17 \times 10^{-6}$ | $2,81 \times 10^{-5}$ |
| *Ae. speltooides* | Ribosome | 44 | 24,91 | 0,21 | 0,06 | $2,21 \times 10^{-6}$ | $2,81 \times 10^{-5}$ |
| *Ae. speltooides* | Pentose phosphate pathway | 12 | 15,21 | 0,29 | 0,07 | $3,06 \times 10^{-6}$ | $3,26 \times 10^{-5}$ |
| *Ae. speltooides* | Nucleotide metabolism | 18 | 18,18 | 0,30 | 0,10 | $3,58 \times 10^{-5}$ | $2,53 \times 10^{-4}$ |
| *Ae. speltooides* | Efferocytosis | 17 | 22,54 | 0,27 | 0,10 | $5,80 \times 10^{-5}$ | $3,35 \times 10^{-4}$ |
| *Ae. speltooides* | Cadherin signaling | 10 | 11,11 | 0,28 | 0,05 | $3,50 \times 10^{-4}$ | $1,27 \times 10^{-3}$ |
| *Ae. speltooides* | Protein export | 13 | 16,62 | 0,28 | 0,09 | $6,21 \times 10^{-4}$ | $2,02 \times 10^{-3}$ |
| *Ae. speltooides* | Carbon fixation by Calvin cycle | 15 | 18,65 | 0,18 | 0,07 | $1,25 \times 10^{-3}$ | $3,19 \times 10^{-3}$ |
| *Ae. speltooides* | Pyrimidine metabolism | 9 | 8,82 | 0,36 | 0,10 | $4,25 \times 10^{-3}$ | $7,59 \times 10^{-3}$ |
| *Ae. mutica* | Protein export | 15 | 15,99 | 0,29 | 0,11 | $1,20 \times 10^{-3}$ | $3,11 \times 10^{-3}$ |

The next steps include fine-mapping to identify candidate adaptive polymorphisms underlying the detected signals, followed by functional validation such as VIGS or CRISPR-Cas9 (W. Wang et al. 2022; Waites et al. 2025) and marker development (Edet et al. 2018). Given the close relatedness of the studied species to the B-genome donor lineage, validated favorable alleles could then be evaluated for introgression into elite wheat backgrounds through approaches such as backcrossing or speed breeding. Ultimately, integrating evolutionary genomics with functional validation and breeding will be key to mobilizing CWR diversity and improving crop resilience.

## 5 Data availability

Processed polymorphism and divergence alignments, and the processed result tables underlying the reported analyzes — including candidate gene lists, GO enrichment results, protein–protein interaction networks, and polymorphism/divergence statistics for each species — are deposited in the Zenodo repository at https://doi.org/10.5281/zenodo.21397724, link: (preview link).

The code used to perform the analyzes can be found in https://github.com/manuelbt-web/Branch_specific_MKT/tree/main.

Supplementary figures and tables are provided in a separate file, Supplementary_material.pdf, submitted along with the main manuscript.

## 6 Acknowledgments

We thank G. Sylvain for helpful discussions and suggestions, and Johanna Girodolle and A. Morganne for their assistance in using Gecko.

